# Transcriptomics of cold stress and recovery reveal strongly tissue-specific responses

**DOI:** 10.64898/2026.05.19.725261

**Authors:** Mara Heilig, Lahari Gadey, Jenna Tomkinson, James A. deMayo, Gregory J. Ragland

## Abstract

Cellular stress responses are often characterized as conserved, cell-autonomous processes. However, it remains unclear whether stress responses are coordinated uniformly across tissues within complex organisms, particularly during ecologically relevant conditions. We investigated tissue- and stage-specific transcriptional responses to cold stress in *Drosophila melanogaster*. Adults and larvae were independently exposed to a gradual cooling and recovery time series, and three adult tissues (gut, ovary, brain) and one larval tissue (gut) were sampled at baseline, at two time points that spanned the critical thermal minimum (before and during chill coma), and after recovery to rearing temperature. Transcriptomic analyses revealed strongly tissue- and stage-specific responses to cold stress, with limited overlap in differentially expressed genes or functional enrichment across tissues. These results indicate that the organismal response to thermal stress at the transcriptional level is not coordinated by a unified transcriptional program, but rather by largely distinct, tissue-specific regulatory processes.

## INTRODUCTION

Organisms across diverse taxa face extreme environmental conditions that disrupt normal cellular and physiological functions and threaten homeostasis (Kültz, 2003; Kültz, 2005; Kültz, 2020a; Schulte, 2014). These hostile environments, characterized by temperatures, salinities, oxygen levels or other ecological factors outside of an organism’s optimal performance range, can elicit physiological stress responses operating from the cellular to organismal level (Miller and Stillman, 2012). In ectotherms, for example, environmental temperature directly influences cellular processes like membrane fluidity and enzymatic activity, which in turn impacts an organism’s performance and fitness. Understanding how organisms respond to these perturbations at various levels of biological organization (i.e. cellular, tissue, organismal), is a fundamental goal in ecological physiology.

To cope with stressors that induce some form of macromolecular damage and disrupt cellular homeostasis, organisms have evolved a conserved set of molecular responses, often termed the cellular stress response (CSR) (Kültz, 2003; Kültz, 2005; Kültz, 2020a). The CSR is primarily characterized by the energy-intensive process of macromolecular repair and stabilization, typically through upregulation of heat shock proteins, molecular chaperones and other protein and DNA repair pathways. By prioritizing cell cycle checkpoints, the CSR helps to minimize the transfer of molecular damage to daughter cells while allowing time for repair and energy repartitioning. These key features have been documented across a wide range of taxa (Kültz, 2020a; Lindquist, 1986; Taylor and Lehmann, 1998), suggesting they have a crucial, evolutionarily conserved role in enabling organisms to respond to extreme environmental perturbations.

Whereas the CSR is well characterized at the cellular level (Fulda et al., 2010; Kültz, 2020b; Kültz, 2020a), it remains unclear whether complex organisms coordinate these responses uniformly across tissues/cell types or through tissue/cell-specific strategies. Multicellular organisms are not simply collections of independent cells that respond uniformly to environmental challenges. Instead, they are composed of specialized cells with distinct functions that can respond to environmental cues and stressors in a tissue-specific manner. In insects, for example, cold exposure elicits ion and water imbalance in the gut leading to increased potassium concentrations in the hemolymph (Des Marteaux et al., 2017; Yerushalmi et al., 2018), while the central nervous system (CNS) undergoes neural depolarization and disrupted ion transport leading to a whole-body chill coma response (Andersen and Overgaard, 2019; MacMillan and Sinclair, 2011). Disruption of ion and osmotic balance in muscle tissue also contributes to immobility in the cold (Findsen et al., 2016; MacMillan and Sinclair, 2011). Reproductive tissues may also respond idiosyncratically to cold stress by blocking oogenesis in the ovaries as a result of egg chamber degeneration (Gandara and Drummond-Barbosa, 2022; Lirakis et al., 2018), leading to reproductive dormancy (Lirakis et al., 2018). Finally, in holometabolous insects, transcriptomic responses to cold can differ markedly across developmental stages (Freda et al., 2022), reflecting tissue-specificity during the reorganization of tissues from juvenile to adult stages. Collectively, these results demonstrate tissue-level, specialized responses to cold. However, the conceptual CSR framework would predict some signal of common cellular responsess across tissues as well.

High-throughput “‘omics” approaches, most prominently transcriptomics, proteomics, and metabolomics, provide powerful tools to test for an *in vivo* conserved CSR across tissue types by generating global snapshots of molecular changes in response to stress. However, many omics-enabled studies homogenize whole organisms, potentially obscuring tissue-specific responses. Certainly, many omics-enabled studies also quantify tissue-specific responses (Fagerberg et al., 2014; Li et al., 2022). But, very few studies compare stress responses across multiple tissues (but see Jin et al., 2022; Zhang et al., 2017), and generally the focus of these studies is on a general understanding of a particular stress response, and not on the tissue specificity of the response *per se*. Here we explore tissue-specificity of whole-transcriptome responses to cold stress in *Drosophila melanogaster* flies across tissue types within and between developmental stages. We suggest three scenarios for how organisms transcriptionally respond to stress across tissues. First, under a convergent CSR model, the same core genes and pathways, such as HSPs and DNA repair pathways, would be activated uniformly across all tissues (Kültz, 2020a). Alternatively, tissues might achieve similar functional outcomes, but through different transcriptional programs. In this scenario, we would predict functional enrichment of processes related to metabolic reprogramming and molecular repair, but through distinct transcriptional profiles. Finally, tissues might exhibit entirely distinct transcriptional and functional profiles, indicating a tissue-specific division of labor where different tissues specialize in distinct processes which may or may not involve canonical CSR mechanisms. We emphasize that these hypotheses could apply differently at different, hierarchical levels of cellular regulation (e.g., transcripts, proteins, and metabolites), and thus inferences from this study are limited to transcriptional patterns.

To determine if a conserved CSR operates across tissues and developmental stages, we examined whole-transcriptome responses to cold stress in three adult tissues (gut, ovary and brain) and one larval tissue (gut) in *D. melanogaster*. We exposed flies to ramping temperatures starting with a baseline of 21°C, decreasing to 0°C, then rapidly warming to a recovery temperature of 21°C. Previous work has shown that cooling down to these temperatures (0°C) elicits a robust transcriptional response in whole body homogenates of adult *D. melanogaster* (Ullah et al., 2024) and that larvae will transcriptionally respond to temperatures as low as -5°C (Freda et al., 2022). The temperature at which locomotion ceases in *D. melanogaster*, also known as the critical thermal minimum (CT_min_) is approximately 5-5.5°C (MacLean et al., 2019; Zarubin et al., 2020). By sampling flies at cold temperatures above and below CT_min_ and at a warmer temperature following cold exposure we captured transcriptional dynamics in response to 1) mild cold stress that causes reduced mobility above CT_min_, 2) moderate cold stress that causes complete loss of neuromuscular function but not immediate death (Andersen et al., 2015) or pronounced performance deficits (Garcia and Teets, 2019) following recovery, and 3) during recovery at warmer temperatures.

## MATERIALS AND METHODS

### Insect Lines and Rearing

We obtained *D. melanogaster* flies of the Oregon R strain from the Bloomington *Drosophila* Stock Center (Bloomington IN, USA) at Indiana University and maintained them at 21°C on 12h:12h light/dark cycle (Percival DR-36VL incubator, Percival Scientific, Perry, IA). All flies were reared on a diet consisting of cornmeal, molasses and yeast, and agar, with propionic acid and Tegosept (Genesee Scientific, Morrisville NC, USA) as a preservative (Recipe in Supplemental Data A).

To initiate experiments, 5 male and 5 female flies were sorted under CO_2_ immediately after eclosion then placed in new food vials. Adults were transferred to new food vials for mating and oviposition every day for 4 days, discarding the vials from the first day to avoid any lingering effects from the CO_2_ used in sorting. Larval offspring were reared to 3^rd^ instar (likely representing a mixture of males and females) and placed into fresh vials for use in experiments. Adult offspring (the generation sampled for the experiments) were collected from the vials established by the parental generation within 1 day of eclosion (12-14 post-oviposition) and sorted by sex. Males were discarded, while females were placed into fresh vials and used in experiments. All adult flies used in experiments were adult virgin females aged 4 days old to avoid any lingering effects of CO_2_ used during sorting.

### Experimental Treatments

To determine how tissue- and development-specific transcriptomic profiles change during a chill coma-inducing cold exposure and then during recovery, we sampled various tissues (see below) from flies before, during, and after a ramping cold exposure. Individual adults or 3^rd^ instar larvae were aspirated (adults) or transferred as described in Freda et al. (Freda et al., 2017) into 1.7ml microfuge tubes (including a small, wetted piece of cotton in the lid to prevent desiccation) that were then inserted into an aluminum block that was cooled and heated through copper coils using a temperature-controlled, recirculating bath (Arctic A40 recirculating bath, ThermoFisher, Denver CO, USA) (Sinclair et al., 2015). We first sampled individuals at the rearing temperature of 21°C, an unstressed control. Then, we decreased the temperature 0.1°C per minute to 7°C, the second sampling point, and continued until 0°C, the third sampling point (Fig. 1). We expected 7°C to induce mild cold stress just above CT_min_ that does not induce chill coma but causes slowed mobility (Andersen et al., 2015), while we expected 0°C, a temperature below CT_min_, to induce a chill coma that is not sufficient to induce mortality (Andersen et al., 2015; Freda et al., 2017). We visually confirmed that all individuals sampled at 7°C and 0°C were moving, and comatose, respectively. Once the minimum temperature of 0°C was reached, the temperature of the bath was increased as fast as the bath could warm for 14 minutes, at which point the temperature reached 21°C. After 30 minutes at a constant 21°C, samples were collected for the fourth and final time point to assess how transcription was affected as the individual recovered from the cold stress.

**Figure 1.**
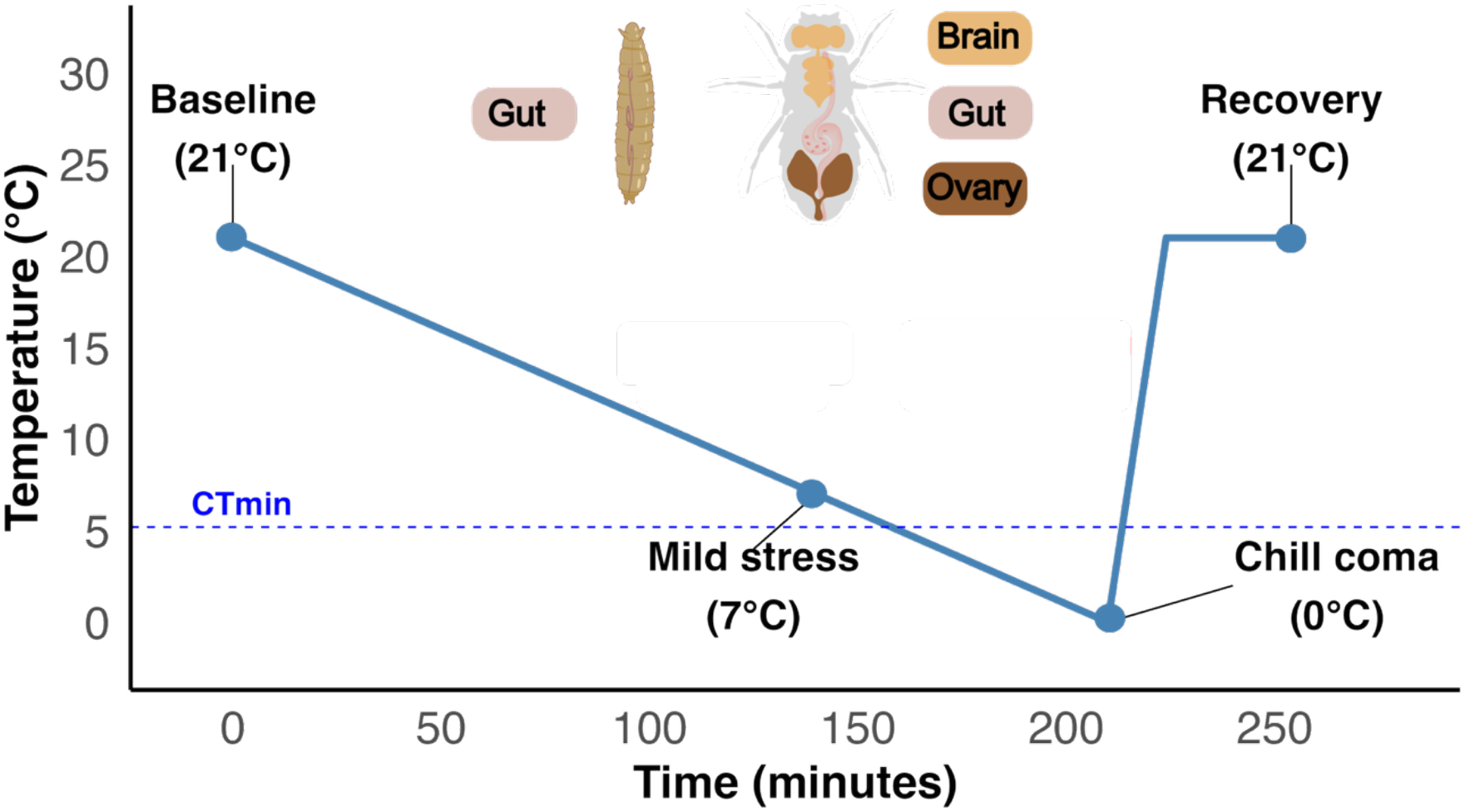
Summary of the experimental temperature profile and sampling scheme for cold exposure in *Drosophila melanogaster*. Temperature was decreased from baseline (21°C) to mild cold stress (7°C) and then to chill coma (0°C) during a controlled cooling phase (0.1°C/min.), followed by rapid warming (14 min.) to the recovery temperature (21°C). Circles denote sampling points at which whole flies were flash-frozen in liquid nitrogen. Adult tissues (brain, gut, ovary) and larval tissue (gut) were dissected from 10-15 pools of flies per biological replicate. A total of 4 biological replicates per tissue per temperature were generated for mRNA sequencing.

### Sampling and RNA extraction

Following each temperature exposure, microcentrifuge tubes with individuals were removed from the block and snap frozen with liquid nitrogen. Tubes were stored at -80°C for 6-10 weeks prior to dissection. Brain, gut and ovary tissues from adults, and gut tissue from larvae were dissected separately into cold RNA*later* (Thermo Fisher, Waltham, MA, USA). Tissue was pooled from 10-15 individuals in TRI reagent (Zymo Research, Irvine CA, USA). Four adult pools per tissue per temperature yielded a total of 48 adult samples with four biological replicates per treatment (4 pools x 3 tissues x 4 temperatures). Four larval gut pools per temperature yielded a total of 16 larval samples (4 pools x 1 tissue x 4 temperatures). RNA from each biological replicate was extracted using Zymo Direct-zol RNA kits (R2051) according to the manufacturer’s instructions, including the optional DNase I treatment to eliminate DNA contamination. RNA purity, quantity and integrity were determined using a NanoDrop 2000 (Thermo Fisher, Waltham, MA, USA) and TapeStation 4200 (Agilent, Santa Clara, CA, USA) analysis.

### Library preparation, sequencing and initial read processing

Library preparation and mRNA sequencing were performed at the University of Colorado Anschutz Medical Campus Genomics Core using the Universal Plus mRNA-seq stranded library preparation kit with NuQuant (TECAN, Männedorf, Switzerland). An input of 200ng total RNA was used to generate 150 bp, paired-end RNA-Seq reads on a NovaSeq X (Illumina) sequencer at a target depth of 20 million paired-end reads per sample. Raw sequencing reads were de-multiplexed using bcl2fastq. Inspection of initial data originating from adult tissues revealed higher than expected levels of ribosomal RNA and relatively low read counts (mean of 15.7 million paired-end reads per sample). To provide adequate coverage depth, each adult tissue library was re-sequenced across two additional flow cells to yield a total sequencing depth of

80.4 million paired-end reads per sample. Libraries prepared from larval guts were created in a different batch that did not suffer from overly high ribosomal RNA contamination and yielded a mean of 33.4 million paired-end reads per sample. Raw sequence reads were archived in the NCBI SRA (PRJNA1456191), and full details of the bioinformatics workflow can be found on the provided Zenodo link for review.

Briefly, raw reads were trimmed to remove adapters using fastp (Chen et al., 2018). Because adult reads were sequenced across 3 flow cells (see above), all trimmed reads for each sample were concatenated into a single file prior to mapping. Larval samples were only sequenced once, and thus none needed to be concatenated. All trimmed reads were mapped to the indexed *D. melanogaster* r6.49 genome using STAR version 2.7.6a and read counts per gene were generated using default parameters, except that multi-mapping reads were excluded (--outFilterMultimapScoreRange 0 and –outFilterMultimapNmax 1), and we enabled merging of overlapping mate pairs with a minimum of 8 base overlap between read pairs (--peOverlapNbasesMin 8) and no more than 20% mismatches in the overlap region (--peOverlapMMp 0.2).

### Statistical modelling of temperature and tissue specific transcription (adults and larvae)

Gene level read counts from the forward strand were filtered to retain only genes with at least one read in 50 percent of samples. Libraries were normalized using the trimmed mean of M-values (TMM) method via the edgeR package in R (Robinson and Oshlack, 2010). We identified the 500 genes showing the greatest variation in expression (separately for adults and larvae) and visualized their relationships using multi-dimensional scaling (MDS) plots (Robinson and Oshlack, 2010). One adult ovary sample at 7°C (14_Ovary_7C) was removed from subsequent analyses due to its exceptionally high normalization factor (2.17), and its appearance as a clear outlier in the adult MDS plot (Fig. S1a). Similarly, one larval gut sample at 0°C (Gut-0-R3) was removed from all subsequent analyses due to its low normalization factor (0.66), outlier status in the larval MDS plot (Fig. S1b), and exceptionally low read depth. Following outlier removal, TMM-normalized count data were converted to log_2_ counts per million (logCPM) with a prior count of 3 to stabilize variance estimates for lowly expressed genes (Law et al., 2014). To account for global differences in expression distributions across adult tissue libraries while preserving biological variation within tissues, we applied quantile smoothing (qsmooth) normalization (Hicks et al., 2018) using tissue as the grouping factor. The qsmooth approach relaxes the assumption that all samples share the same distribution and computes a weight at each quantile by comparing variability between tissues relative to within tissues. We did not apply qsmooth normalization to larval libraries because all samples were from a single tissue.

**Figure S1.**
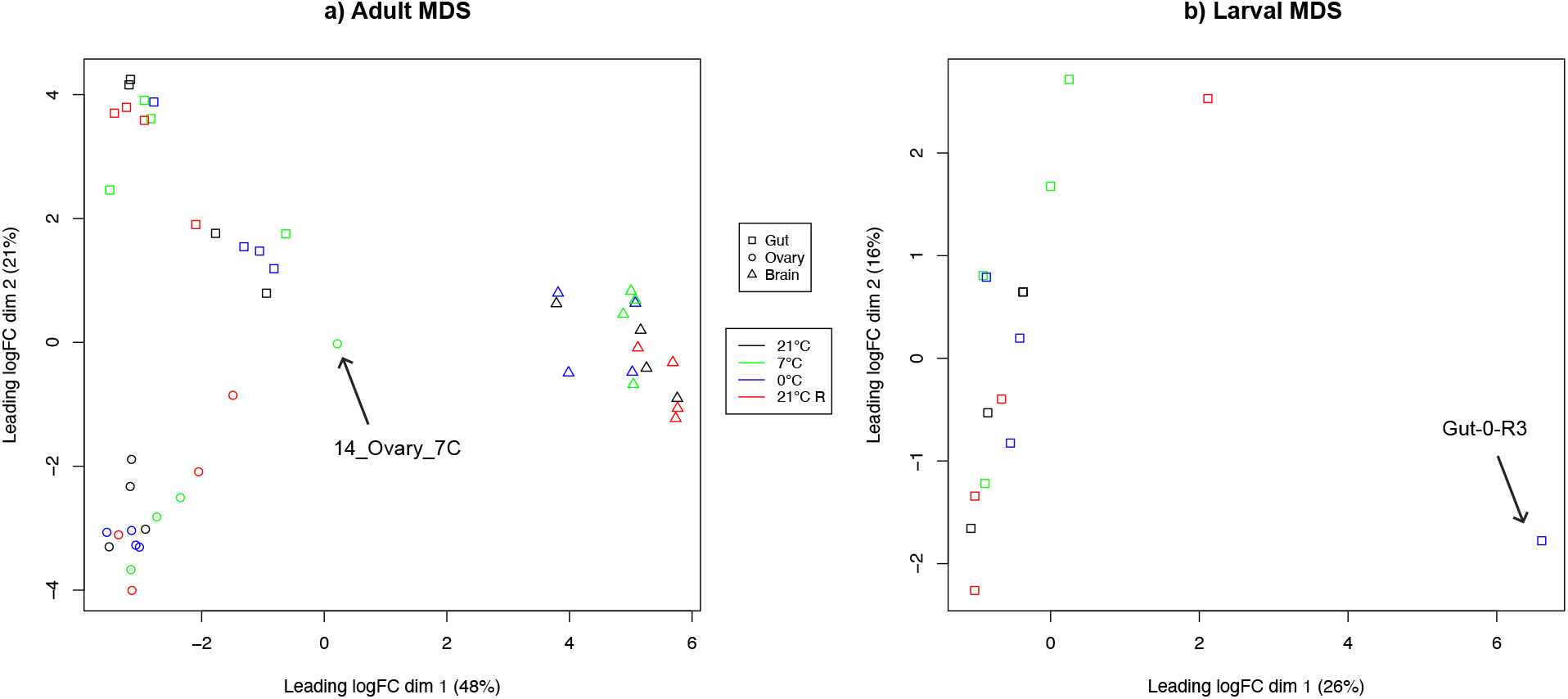
Multidimensional Scaling (MDS) analysis of the 500 most differentially abundant transcripts in (a) adults and (b) larvae following cold exposure (7°C and 0°C) and recovery (21°C). Each RNA library (sample) is plotted with shape indicating tissue and color indicating sampling temperature. The sample ID for outliers that were removed from all analyses are indicated with arrows pointing to the sample.

Differential gene expression was assessed separately for adult and larval libraries using limma-trend (Ritchie et al., 2015). For adults, we fit linear models to qsmooth-normalized logCPM values with tissue (three levels; brain, gut, and ovary), temperature treatment (four levels; 21°C as reference, 7°C, 0°C, 21°C recovery), and their interaction as fixed effects (gene expression ∼Tissue*Temperature). For larvae, we fit linear models to logCPM values with temperature as the sole fixed factor (gene expression ∼Temperature). Coefficients for both adult and larval models were estimated using lmFit, and empirical Bayes moderation with a mean-variance trend was applied. We also performed all pairwise temperature contrasts within each tissue (including larval gut) using custom contrast matrices. For each coefficient and contrast, differentially expressed (DE) genes were identified at a false discovery rate (FDR) < 0.05 using the Benjamini-Hochberg (BH) correction. Genes showing differential expression (FDR < 0.05) in at least one pairwise contrast within a tissue were classified as temperature-responsive in that tissue. Log_2_ fold changes and *p*-values were extracted to identify tissue-specific vs. overlapping (across tissues) DE genes, and to construct gene expression trajectories across the temperature time course for downstream enrichment analyses.

### Enrichment of shared DE genes

To assess whether DE genes overlapped more than expected by chance across tissue and stage comparisons, we performed Fisher’s exact tests on pairwise contingency tables and estimated odds ratios and associated 95% confidence intervals (CIs). For adult tissue comparisons (gut vs. ovary, gut vs. brain and ovary vs. brain), genes were classified as DE if they had an FDR < 0.05 in at least one temperature contrast. For the cross-stage comparison (adult gut vs. larval gut), we used the union of all genes detected in both datasets as the background because adult and larval samples were analyzed separately and had different sets of expressed genes. Genes were classified as DE in adult and larval guts if they showed differential expression (FDR < 0.05) in at least one temperature contrast within the tissue.

### Functional enrichment analysis

We used the DAVID functional annotation clustering tool from NCBI (Huang et al., 2009) to identify functional categories enriched in the sets of transcripts illustrating temperature-specific responses within tissue. Functional categories included Uniprot keyword searches (UPK), Gene Ontology groups (GO), Interpro protein domains (INTERPRO) and Kyoto Encyclopedia of Genes and Genomes pathways (KEGG). Enrichments were analyzed separately for genes that were up- and down-regulated at each temperature comparison within each tissue. We considered an enrichment score >2.0 and FDR<0.05 as evidence for enrichment of a functional category. To illustrate overall functional categories, we filtered specific terms into broader, representative categories (see Supplemental Data L for category filtering).

### RESULTS

#### Sets of differentially expressed genes are primarily tissue- and stage-specific

We first assessed the overall effects of tissue, temperature, and their interaction on gene expression in adult tissues. Global transcriptional patterns were primarily driven by tissue-specific expression, with 10,532 genes displaying significant differences among brain, gut and ovary in adults (FDR < 0.05). Temperature had minimal effects independent of tissue with only 3 genes similarly differentially expressed across temperatures, while 313 genes showed a significant temperature by tissue interaction. Using pairwise contrasts, we also identified transcripts that were DE across temperatures within tissues. A total of 1043, 3359, 191 and 9 genes were DE in at least one temperature comparison in larval gut, adult gut, adult ovary and adult brain, respectively (Fig. 2). Thus, adult and larval gut transcription were highly temperature-responsive, ovary transcription was moderately responsive, and brain transcription was nearly unresponsive.

**Figure 2.**
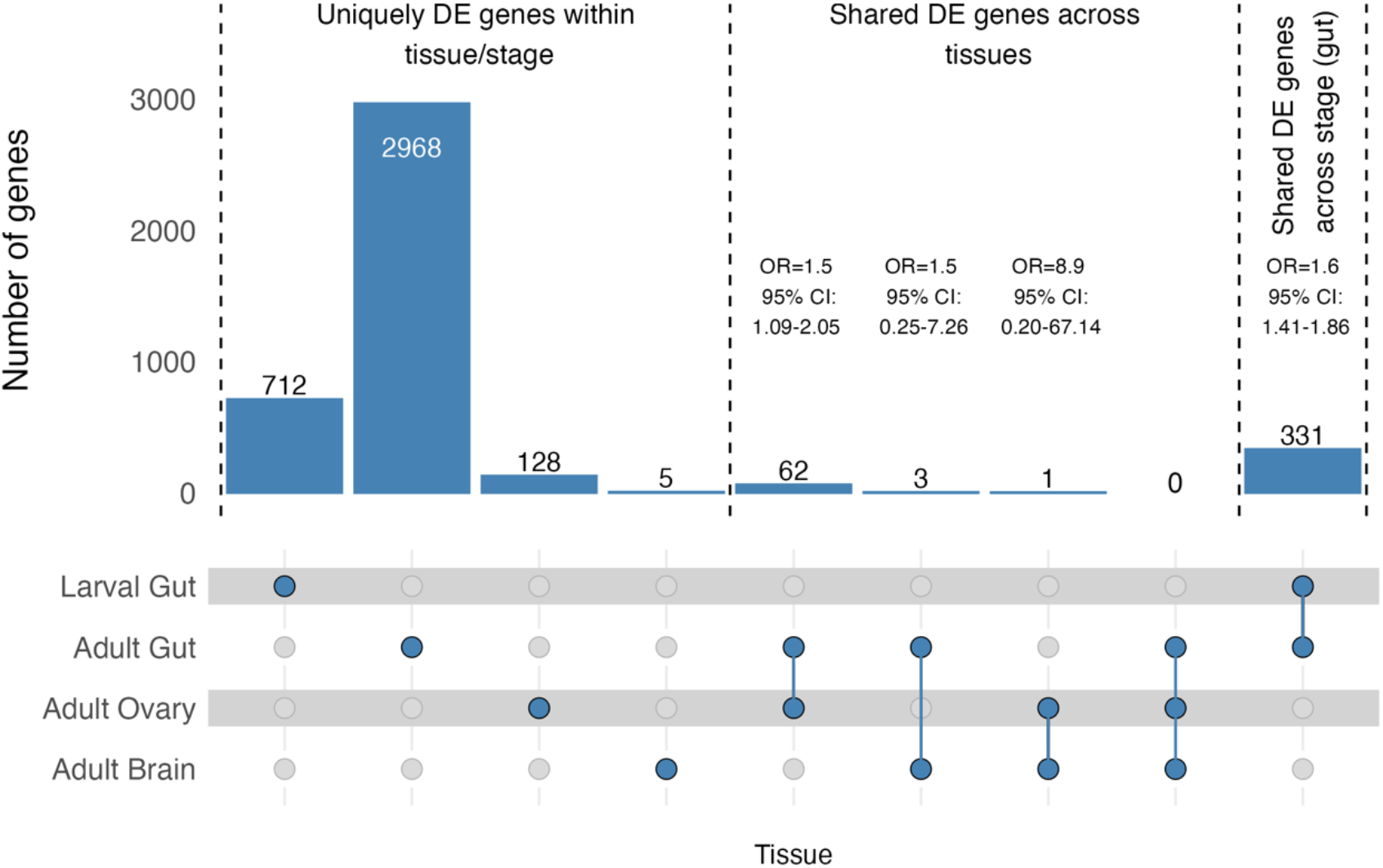
Differentially expressed genes are primarily tissue- and stage-specific. Bars show the number of genes that are uniquely DE within a tissue/stage or overlapping across defined combinations. Adult-only overlaps (shared DE genes across tissues) are shown separately from cross-stage (adult vs. larval) gut overlaps. Because our inference is limited to across tissues within stage (adults) and across-stage within tissue (guts), we did not evaluate larval gut vs adult ovary/brain. For pairwise comparisons, bars are annotated with estimated odds ratios (OR) and 95% confidence intervals (CI). OR > 1 suggests higher overlap than expected by chance, with CI width reflecting uncertainty).

Limited overlap in the sets of temperature-responsive (DE) genes among tissues and across stages between any temperature support responses to cold that were largely tissue- and stage-specific. Though there was moderate enrichment of shared DE genes between adult gut and ovary, there was no more overlap in DE genes in other adult tissue comparisons than expected by chance (Fig. 2). There was also moderate enrichment of shared DE genes between adult and larval gut, though not more than between adult gut and ovary (95% CI of OR estimates overlap). In all comparisons, however, most DE genes were tissue- and stage-specific.

### Differentially expressed transcripts are inconsistently correlated between tissues and stage

Gene expression changes for transcripts DE across consecutive time/temperature comparisons tended to be DE in the same direction in the adult gut and adult brain (Fig 3g-i), but in opposite directions in the adult gut and adult ovary (Fig. 3d-f). Transcripts that were DE from 21°C to 7°C and 7°C to 0°C tended to be slightly negatively correlated in the adult-gut and larval-gut (Fig 4a-b). Conversely, transcripts that were DE from 0°C to the recovery temperature (21R) tended to be positively correlated (Fig. 3c). Thus, when considering a more inclusive set of genes (DE in either tissue in a pairwise comparison) there was a statistical signature of relatedness among tissue/stage transcriptomic responses to cold. However, the frequently observed patterns of anticorrelation suggest that tissues in general may exhibit unique responses even for genes that may be responsive across more than one tissue.

**Figure 3.**
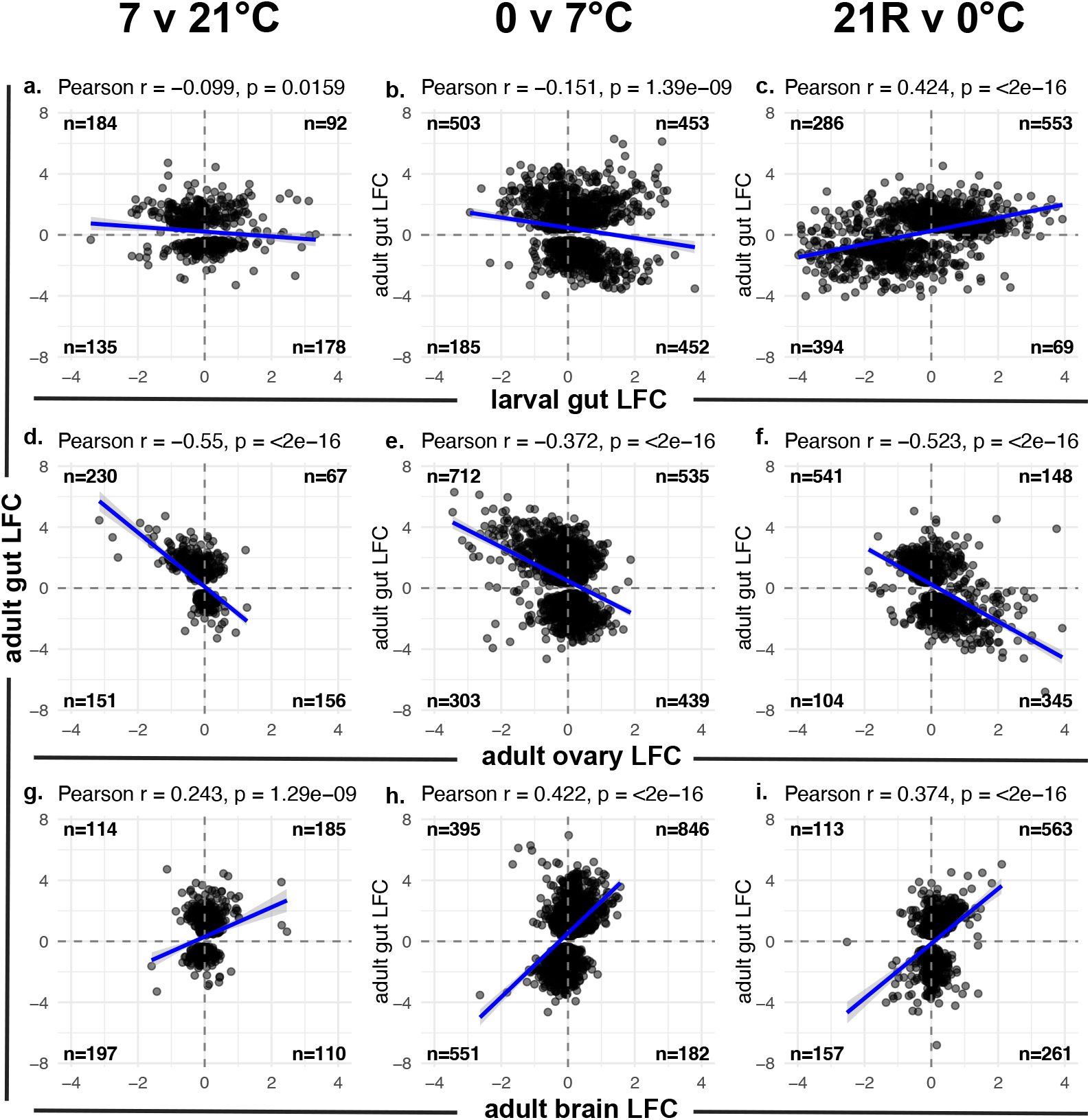
Mixed correlation patterns of log_2_ fold-change (LFC) responses across tissues and stages for transcripts DE in pairwise comparisons across three temperature contrasts. Each quad-plot shows LFCs for genes DE in either or both tissues for a given temperature comparison. Columns represent temperature transitions: 21°C to 7°C (left) 7°C to 0°C (center) and 0°C to 21°C at recovery (right). Rows correspond to the relationship between the LFC of genes in adult gut (y-axis) and the contrasted tissue/stage (x-axis): larval gut (a-c), adult ovary (d-f) and adult brain (g-i). The strength and direction of relationships were evaluated using Pearson correlation coefficients.

**Figure 4.**
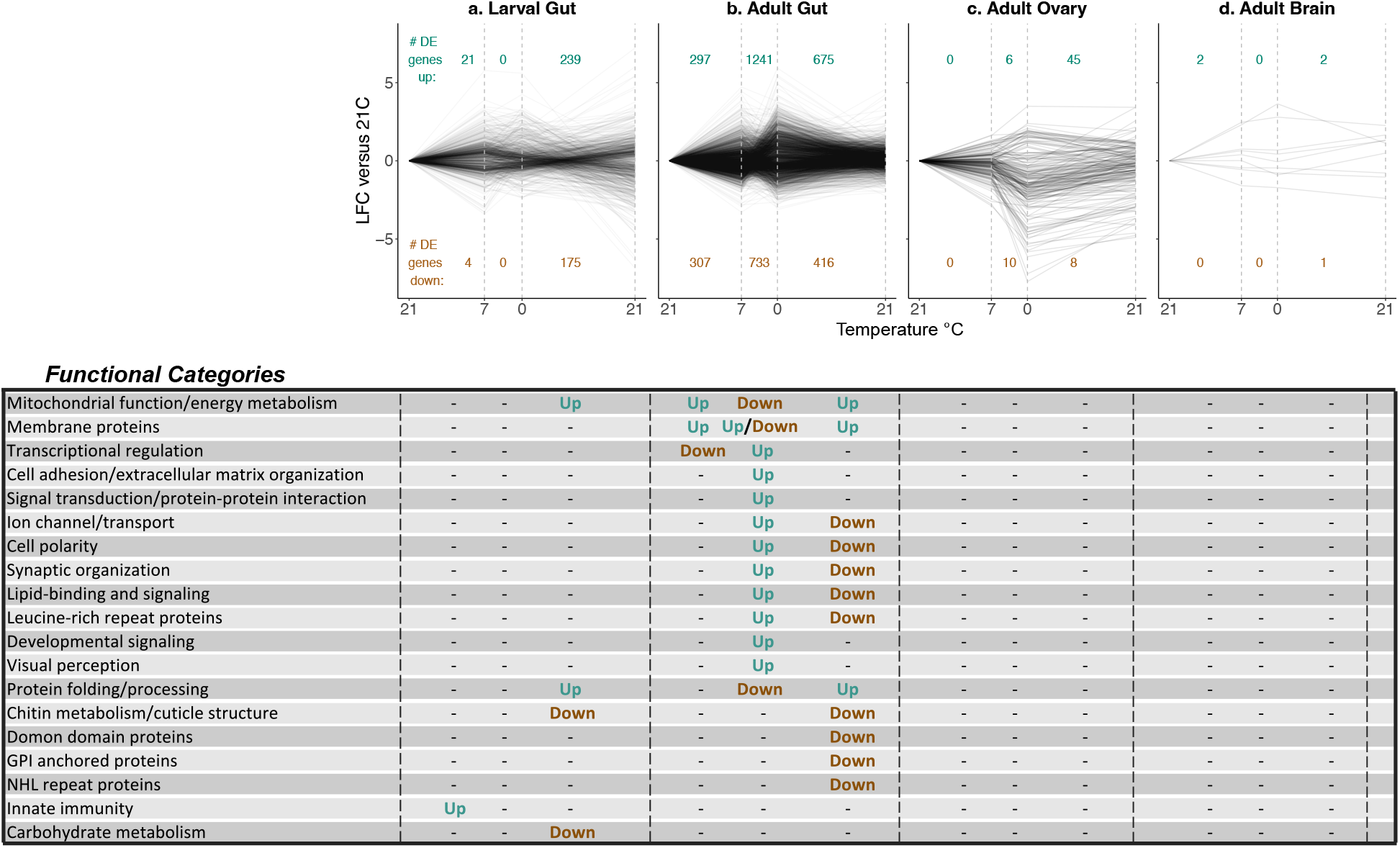
Trajectories of differential expression (DE) during and after cold exposure for DE transcripts (FDR<0.05) across any temperature contrast. **Panels correspond to** a) Larval Gut, b) Adult Gut, c) Adult Ovary, d) Adult brain. The number of up- and down-regulated genes corresponding to each consecutive temperature transition are shown within each trajectory. Below trajectories, functional categories that were enriched for genes that were up or down regulated are shown below their respective temperature transitions. Dashes indicate no enrichment.

### Distinct trajectories and minimal overlap of functional enrichment across tissue and stage

For each tissue and stage (adult-gut, adult-ovary, adult-brain and larval-gut), we plotted the log_2_ fold change relative to 21°C across consecutive temperatures (21°C vs. 7°C, 7°C vs. 0°C, 0°C vs. 21°C recovery) for any gene that was differentially expressed between any temperature comparison (Fig. 4a). Overall, DE genes showed distinct trajectories across temperatures within each tissue and stage. Further, most DE across consecutive temperature/timepoints in both larval and adult guts occurred during the recovery period (from 0°C back up to 21°C), whereas most DE in adult ovaries occurred during the ramp from 21°C to 0°C. The adult brain had very few DE genes across any consecutive timepoint.

Among genes DE at any temperature contrast, the overall direction of gene expression change differed pre- and post-CT_min_ in larval and adult gut (Fig. S2a-b) but remained positively correlated in adult ovary (Fig. S2c). The magnitude of expression change also differed pre- and post CT_min_ in a tissue-specific manner. In adult gut, mean absolute log_2_ FC increased from 0.65 ± 0.02 for the 7°C vs 21°C comparison to 1.15 ± 0.03 for the 0°C vs 7°C comparison. A similar pattern was observed in adult ovary (7°C vs 21°C: 0.62 ± 0.08; 0°C vs 7°C: 1.35 ± 0.15) whereas larval gut exhibited a more modest decrease in the magnitude of expression change (7°C vs 21°C: 0.74 ± 0.04; 0°C vs 7°C: 0.55 ± 0.03).

**Figure S2.**
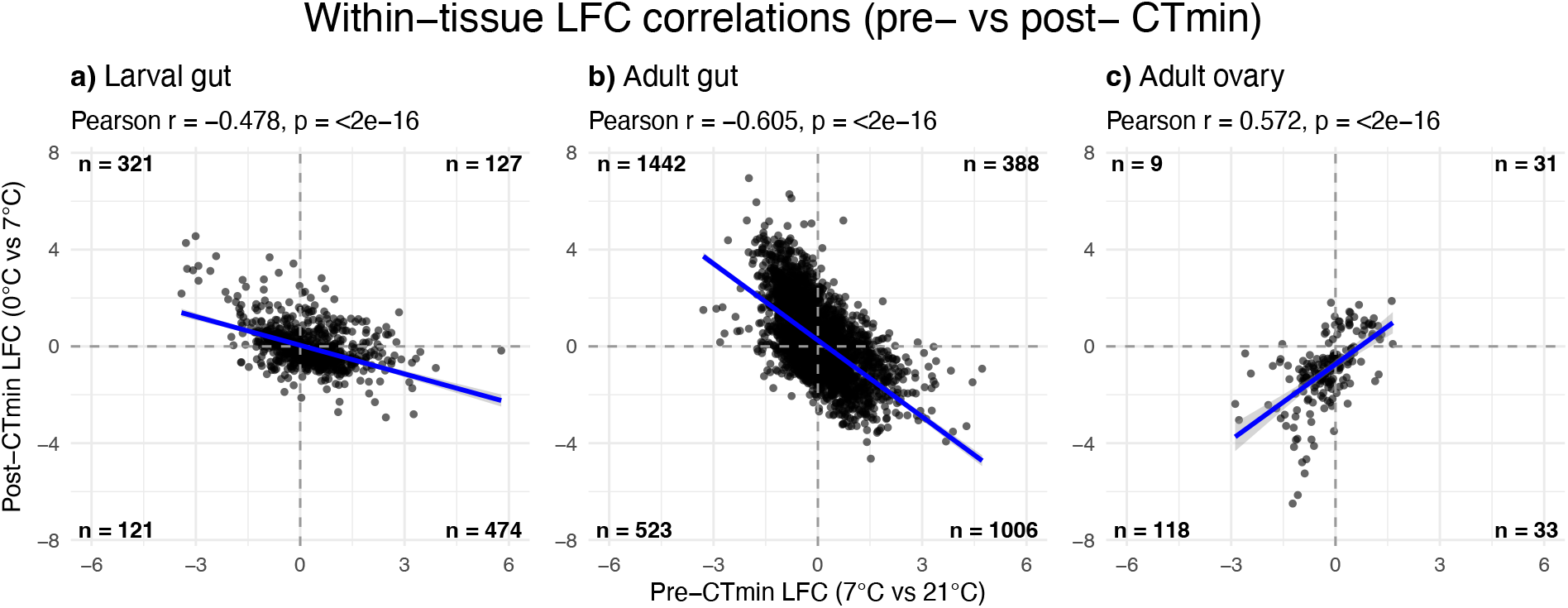
Mixed correlation patterns of log_2_ fold-change (LFC) responses within tissues across CTmin. Each quad-plot shows LFCs for the temperature transition just above (x-axis, 21°C vs 7°C) and below (y-axis, 7°C vs 0°C) CTmin for any gene DE at any time point within that tissue. Pearson correlation coefficients indicate the strength and direction of the relationship pre- and post-CTmin expression changes. Adult brain is not shown due to insufficient DE genes for correlation analysis.

The full results of enrichment analysis conducted with the David functional annotation tool (Huang et al., 2009) can be found in Supplemental Data C-K. Cluster analysis identified functional categories and terms that cluster together based on gene overlap; we then designated a collective term for any cluster with an enrichment score > 2 based on the annotation term of categories with a Benjamini-Hochberg corrected *p*-value <0.05. Because some collective terms had overlapping functional annotations, we combined collective terms into global terms, which are presented in the bottom panel of Figure 4.

Overall, enrichment analysis revealed minimal functional overlap of DE genes across tissues/stages. The tissues with the most DE genes (adult and larval gut) predictably yielded more enriched clusters due to higher statistical power. Across stages within tissue (gut), there were three functional categories shared (and with the same directionality of DE), but only during the recovery phase. These included mitochondrial function/energy metabolism, protein folding/processing, and chitin metabolism/cuticle structure. However, none of the functional categories enriched in the adult gut cold response were enriched in ovary and there were too few genes DE in brain to meaningfully assess functional enrichment in that tissue (Fig. 4, Supplemental Data C-K).

## Discussion

Understanding how organisms coordinate stress responses across specialized tissues remains a knowledge gap in ecological physiology. We examined immediate transcriptional responses to cold stress and recovery across multiple tissues and developmental stages in *D. melanogaster*. While previous studies have characterized whole-organism or single-tissue responses (see studies cited in “Genomics and transcriptomics” section of (Ullah et al., 2024)), very few studies have compared tissue-specific transcriptomic dynamics throughout the same stress event in any organism (but see Huang et al., 2025 in fish) and none that we are aware of in insects. Our experimental design captures transcriptional profiles across distinct temperature transitions throughout cooling and recovery, supporting a rapid cold stress response that is characterized by tissue- and stage-specific transcriptional and functional profiles.

### Signatures of transcription during cooling

Our results demonstrate that adult and larval *D. melanogaster* tissues exhibit a transcriptional response down to temperatures near or below the induction of chill coma. Though some studies of whole organism homogenates in other insect species have suggested limited transcriptional responses at temperatures near or below CT_min_ (Sinclair et al., 2007; Teets et al., 2012), others suggest at least a moderate response (e.g., hundreds of transcripts DE at 5°C (Zhang et al., 2024). During cooling down to 0°C, whole body homogenates of adult *D. melanogaster* have a robust transcriptional response (Ullah et al., 2024) and larvae transcriptionally respond to temperatures as low as -5°C (Freda et al., 2022).

Across discrete temperature transitions, transcriptional responses revealed tissue-specific shifts in both the direction and magnitude of gene expression changes around CT_min_. In both adult and larval gut, expression changes below CT_min_ (0°C vs 7°C) tended to oppose those observed during milder cooling (7°C vs 21°C), indicating a reversal of transcriptional trajectories once flies entered chill coma. In contrast, ovarian transcriptional responses below CT_min_ generally amplified changes observed during mild cooling. These results suggest that CT_min_ may act as a threshold for gene regulation, though in a tissue-specific manner.

### Substantial variation in transcriptional plasticity across tissues and developmental stage

Tissues varied substantially in the magnitude of their transcriptomic response to cold, revealing unique thermal response strategies. The adult gut exhibited the greatest transcriptional responsiveness with over ten times the number of DE genes than the ovary and over 100 times the number of DE genes than the brain. That the brain shows the least responsiveness despite its central role in the chill coma response is particularly striking. Cold exposure causes neural depolarization and disrupted ion transport in the CNS, leading to whole-body immobilization which characterizes the chill coma response (Andersen and Overgaard, 2019; Robertson, 2004). Several factors might account for the lack of a strong transcriptional signal in the brain. First, the brain may rely more on rapid post-transcriptional modifications to respond to immediate perturbations, though direct evidence for this is lacking. However, previous studies have shown post-transcriptional modifications contribute to environmental stress responses (Hernández-Elvira and Sunnerhagen, 2022) and can occur rapidly in the brain (Bhat et al., 2022). Second, protective repair mechanisms can be constitutively upregulated in anticipation of stressful events (Barshis et al., 2013) and may contribute to “frontloading” of transcripts in the brain. And third, the rapid time course in our experiments might be insufficient to capture brain-related transcriptional responses. Acclimation responses do occur during cooling at 0.1°C/min ramping rates – ramping at 0.1°C/min lowers CT_min_ by ∼2.5°C compared to cooling at 1.0°C/min in *D. melanogaster* (Kelty and Lee, Jr, 1999). Thus, post-transcriptional modifications or other regulatory mechanisms adjusting the proteome or metabolome must occur during cooling. Consistent with the negligible transcriptomic response in the *D. melanogaster* CNS, transcriptional responsiveness in the brain of carp was also 4-10 fold lower than other tissues (liver, kidney, gill) following a 24-hour ramping cold challenge (Huang et al., 2025). This suggests that attenuated transcriptional responses to cold in the brain may be a more general phenomenon, though further work comparing tissue-specific responses across shorter and longer time series in different species is needed to evaluate this pattern more broadly.

It is also possible that a higher diversity of cell types in the gut and ovary contribute to stronger transcriptional signals compared to the brain. Both the gut and the ovary are composed of a mix of epithelial, muscle and stem cells (Hung et al., 2020; Rust et al., 2020), whereas the dissected brain tissue is predominantly composed of neurons and glia (Davie et al., 2018; Yildirim et al., 2019). However, snRNA-seq in the brain, gut and ovary also demonstrate substantial cell type diversity based on transcriptomic clustering (Hung et al., 2020; Lee et al., 2025; Li et al., 2022). Therefore, the nearly 375-fold difference in responsiveness (number of DE genes) between the gut and brain likely indicates fundamentally different responses to cold stress that are likely at least in part due to functional specialization in addition to cell type composition. Though no studies have compared transcriptome-wide responses to acute thermal stress in insects across the specific tissue types examined in our study, previous work in plants (Zhi et al., 2024), and fish (Huang et al., 2025) also demonstrate unique transcriptional signatures across tissue in response to different stressors. Thus, the transcriptional variation known to exist between tissues under unstressed, baseline conditions (Li et al., 2022) likely contributes to tissue-specific responses to different stressors and exposures times in *D. melanogaster*, as well as other taxa.

Comparing larval and adult gut responses revealed a stronger signal of a shared cold response (in terms of number of DE genes) within tissue across life cycle stage than comparing between tissues within life cycle. However, adult and larval guts also exhibited distinct temporal patterns of differential expression when accounting for temporal patterns across temperature transitions. For example, the larval gut was most responsive during recovery (0°C to 21°C), while the adult gut was most responsive during the transition from moderate to more severe cold (7°C to 0°C). Thus, the overwhelmingly unique transcriptional trajectories supports the interpretation that cold responsiveness is functionally decoupled across life stages and may have evolved independently in *D. melanogaster* (Collet and Fellous, 2019; Freda et al., 2017; Freda et al., 2019; Freda et al., 2022; Gilchrist et al., 1997; Loeschcke and Krebs, 1996; Moran, 1994; Tucić, 1979).

### Correlation patterns reveal mixed directionality of tissue and development transcriptomic responses to cold

The direction of expression changes among DE genes reveals that tissues not only differ in how many genes respond to cold but in whether those genes are regulated in the same or opposing directions. While genes DE in either the adult gut and/or brain tended to change in the same direction across temperature transitions, suggesting a potential coordination between these tissues, genes DE in either adult gut and/or ovary were consistently negatively correlated across temperature transitions, indicating they respond to the same cold signals by regulating transcripts in opposite directions. This anticorrelation suggests that the gut and ovary are not simply eliciting independent responses but instead may have opposing physiological priorities. Though we did not identify strong signals of functional enrichment for anti-correlated genes, the distinct functional roles of these tissues could underlie opposing transcriptional priorities (Li et al., 2022). Additionally, the anti-correlation could reflect a resource allocation trade-off in which cold stress causes a conflict between the energetic demands of somatic survival and reproduction. However, this interpretation remains speculative, and the mechanistic basis of this anti-correlation has yet to be established.

The relationship between larval and adult gut transcriptional responses was more nuanced, with directionality shifting across temperature transitions. Genes that were DE in either the adult and/or the larval gut were slightly negatively correlated during cooling but positively correlated during recovery. The weak negative correlation during cooling supports some degree of a decoupled cold response across life stages (Dierks et al., 2012; Freda et al., 2017; Freda et al., 2019; Herrig et al., 2021; Loeschcke and Krebs, 1996), though it may also reflect differences in adult versus larval function (Aghajanian et al., 2016). Conversely, recovery may engage a more conserved set of transcripts, implying shared repair mechanisms upon return to non-stressful conditions.

### Functional enrichment is largely tissue-specific with partial CSR signal in gut

Because the gut exhibited by far the most substantial transcriptional response to cold, and no functional enrichment was identified for genes DE at consecutive temperature transitions in either the ovary or brain, we discuss functional enrichment within the gut in relation to the canonical cellular stress response (CSR) (Kültz, 2020a; Kültz, 2020b; Somero, 2020). In both the larval and adult gut, we found some evidence of functional enrichment consistent with the CSR. Genes associated with energy metabolism (i.e. related to oxidative phosphorylation and the TCA cycle) were enriched across all consecutive temperature transitions in the adult gut, as well during recovery in the larval gut, suggesting shifts in metabolic investment during these phases. Categories related to protein folding and processing were also enriched among genes upregulated during recovery in the adult and larval gut, consistent with prioritizing protein repair following macromolecular damage, a hallmark of the CSR. Further, ion channel and transport processes were also enriched among genes upregulated during cooling (7°C to 0°C) and downregulated during recovery (0°C to 21°C), aligning with known changes in gut ion regulation during cold stress in insects (Des Marteaux et al., 2017; Yerushalmi et al., 2018).

Several functional categories were also enriched in the adult gut with no clear links to the canonical CSR including cell polarity, developmental signaling and visual perception. Whether these categories reflect tissue-specific aspects of gut physiology that are responsive to cold, such as preparing for cell turnover in anticipation of damage, or represent incidental transcriptional changes remains to be determined. Notably, we detected no enrichment of heat shock proteins (HSPs) in any tissue, despite HSPs being a hallmark of both heat and cold stress (Colinet et al., 2010; Feder et al., 1996; Feder et al., 1997; Rinehart et al., 2007). This may reflect the more rapid nature of our experimental time course, as previous work indicates longer exposure times and more severe cold (0°C) lead to increased expression of some *Hsp* genes, particularly during recovery (Colinet et al., 2010).

### Conclusion: cold stress response is tissue-specific with inconsistent signals of shared responses

Collectively, our results support a model in which tissue identity and developmental stage are the primary drivers of transcriptomic responses to rapid cold stress. While some elements of the CSR appear in the gut (i.e. energy metabolism, protein folding, ion transport), these signals are largely absent from the brain and ovary. The distinct transcriptomic trajectories across tissue and stage, and the anticorrelated expression patterns between adult gut and ovary further highlight that different tissues prioritize different physiological processes in response to the same thermal challenge. Altogether, we detected no consistently strong signal of a conserved cellular stress response across tissues following a rapid cold challenge in *D. melanogaster*

## ACKNOWLEDGEMENTS

This work was supported by high performance compute infrastructure in Research Computing at the University of Colorado, Boulder, a CU Denver Undergraduate Research & Creative Activities award to JT, and NSF grant IOS 2045263 to GJR.

## Notes

**Funding:** This work was supported by a CU Denver Undergraduate Research & Creative Activities award to JT, and NSF grant IOS 2045263 to GJR.

**Data accessibility:** All raw transcriptomic datasets generated in this study have been deposited in NCBI BioProject under accession number PRJNA1456191. All code and additional data not included in the supplementary materials are available on Zenodo and will be publicly accessible upon acceptance of this manuscript.

**Conflict of interest:** The authors declare no competing interests.

### Competing Interest Statement

The authors have declared no competing interest.

